# Predicting Negative Control Drugs to Support Research in Drug Safety

**DOI:** 10.1101/380832

**Authors:** Yun Hao, Nicholas P. Tatonetti

## Abstract

The lack of high-quality reference data is a major limitation in drug safety and drug discovery science. Unreliable standards prohibit the use of supervised learning methods and make evaluation of algorithms difficult. While some data is available for positive examples (e.g. which drugs are associated with a side effect), there are no systematic resources of negative controls. To solve this issue, we introduced SIDERctrl, a computational method that ranks drugs based on the likelihood of not causing a side effect. We applied SIDERctrl to predict negative controls from unreported drugs of 890 side effects in SIDER. Our predictions decreased the false negative rate by one-third according to a validation study using AEOLUS data. Three sets of predicted negative controls by different thresholds of precision were provided, and can be accessed at http://tatonettilab.org/resources/negative-drugs.html. This new reference standard will be important in chemical biology, drug development, and pharmacoepidemiology.

**KEY POINTS:**

- The lack of systematic resources providing negative control drugs limits the performance of existing research in drug safety.
- We developed a novel method that integrated chemical and biological properties a drug and the target proteins to calculate the likelihood of the drug being negative control.
- We applied our method to 890 side effects, and showed that our method significantly decreased the false negative rate of predictions.

## 1. INTRODUCTION

Side effects (SEs) of drugs, defined as any untoward medical occurrence during the administration of pharmaceutical products, are a worldwide public health concern. According to US department of Health and Human Services, SEs account for approximately one third of all hospital adverse events and affect about two million hospital stays annually, prolonging each stay by 1.7 to 4.6 days [1]. Serious SEs cause about 100,000 deaths per year, which has become the fourth leading cause of death in the US [2]. Thus, comprehensive evaluation of SEs is needed for every drug to reduce healthcare costs and improve outcomes.

A traditional way to detect SEs is conducting clinical trials of pharmaceutical products. However, the inherent limitations of clinical trials, such as limited duration and population of study often lead to new SEs being discovered in the post-marketing surveillance [3, 4]. The US Food and Drug Administration (FDA) monitors post-marketing usage of drugs through the Adverse Event Reporting System (FAERS), which receives reports from healthcare providers, patients, and pharmaceutical companies. Banda *et al* developed a new resource named AEOLUS, which provided a curated and standardized version of reports from FAERS by removing duplicate records and applying standardized vocabularies to drugs and SEs [5]. Similarly, Kuhn *et al* created SIDER that contained data on 1,430 drugs, 5,880 SEs and 140,064 pairs of relationships between them by mining the product labels of FDA-approved drugs [6]. Such resources can serve as examples to study the mechanism of SEs, as well as to predict the occurrence of SEs. Previous studies have used the resources to build a variety of supervised learning models that incorporated features such as structural properties of drugs [7], genomic-scale metabolic models [8], and drug-induced gene expression [9], to predict SEs of drugs. When building the reference set of each SE, all these approaches simply regarded drugs that were reported with the SE in FAERS or SIDER as positive standards and all the other unreported drugs as negative standards. However, such method of classification is limited by the data incompleteness and inconsistency among these resources as a significant number of SEs have not been labeled or reported [10]. This will inevitably bring in false negatives in the reference drug set, thus decrease the precision of the model. As a result, previous models showed only moderate performance with median AUROC between 0.6 and 0.65 [7–9]. To solve this issue, we presented SIDERctrl, a computational method that ranks unreported drugs by the likelihood of not causing a side effect. We hypothesized that unreported drugs that share similar properties with positive drugs are more likely to cause the same SE while unreported drugs that do not are more likely to be negative controls. Based on the hypothesis, we built a supervised learning model that incorporates multiple pharmacological features such as ATC code, indications, compound structure, etc. We applied SIDERctrl to the SIDER data consisting 890 SEs and predicted three sets of negative control drugs with a minimum precision of 0.15, 0.26, and 0.37. We validated our predictions using the defined relationships between drugs and SEs from an independent resource AEOLUS, and showed that the predicted negative controls exhibit a low false negative rate close to 6%. Our results provide a high-quality reference set of 890 SEs, which can support future methodological research in drug safety.

## 2. METHODS

### 2.1 Construct the training set for SIDERctrl model

Given a list of drugs that were reported with a SE (such drugs are referred to as “true positive drugs” below), the task of SIDERctrl is to learn a classifier which can differentiate the unreported drugs that are more likely to cause the same SE (referred to as “others” below) from the unreported drugs that are less likely to cause the SE (referred to as “negative controls” below). The training set of SIDERctrl model was derived from combining four reference sets manually curated by a previous study [11], which contained 1,824 relationships between 456 drugs and four SEs. The description of reference sets was detailed in Supplementary Methods. Drugs without target annotation from DrugBank [12] or SE annotation from SIDER [6] were removed.

### 2.2 Calculate pairwise similarity of seven pharmacological features in SIDERctrl model

A similarity score between 0 and 1 was assigned to each drug pair regarding the following features: 1) ATC code: a discrete value from [0, 0.2, 0.4, 0.6, 0.8, 1] based on accumulative similarity in 5 levels of ATC code (first digit, first three digits, first four digits, first five digits, all seven digits); 2) Indication: Jaccard similarity based on annotation from MEDI [13]; 3) Structure: Tanimoto similarity based on SMILES representation; 4) Target: Jaccard similarity based on annotations from DrugBank [12]; 5) Target PPI: proportion of target pairs that share protein-protein interactions based on annotations from BioGRID [14]; 6) Target phenotype: proportion of target pairs that share genetic variant-disease links based on annotations from OMIM [15]; 7) Target sequence: average pairwise sequence similarity between target proteins calculated by R package protr [16].

### 2.3 Build the SIDERctrl model and evaluate the performance

Using the seven features above, each unreported drug was scored by the maximum feature similarity to all true positive drugs (Fig. 1A). The SIDERctrl model contains 100 random forest classifiers [17] built using the training data. The number of trees was set as 500 in each classifier. The results were averaged across 100 classifiers to account for the stochastic nature of random forest. Bootstrap sampling was used to estimate the 95% confidence interval of each measurement. The out-of-bag probability was used to evaluate the performance of the classifier, which was measured by the area under receiver operating characteristic curve (AUROC). The predictive power of each feature was measured by the increase in mean squared error (MSE) when the feature was removed from the model.

**Fig. 1:**
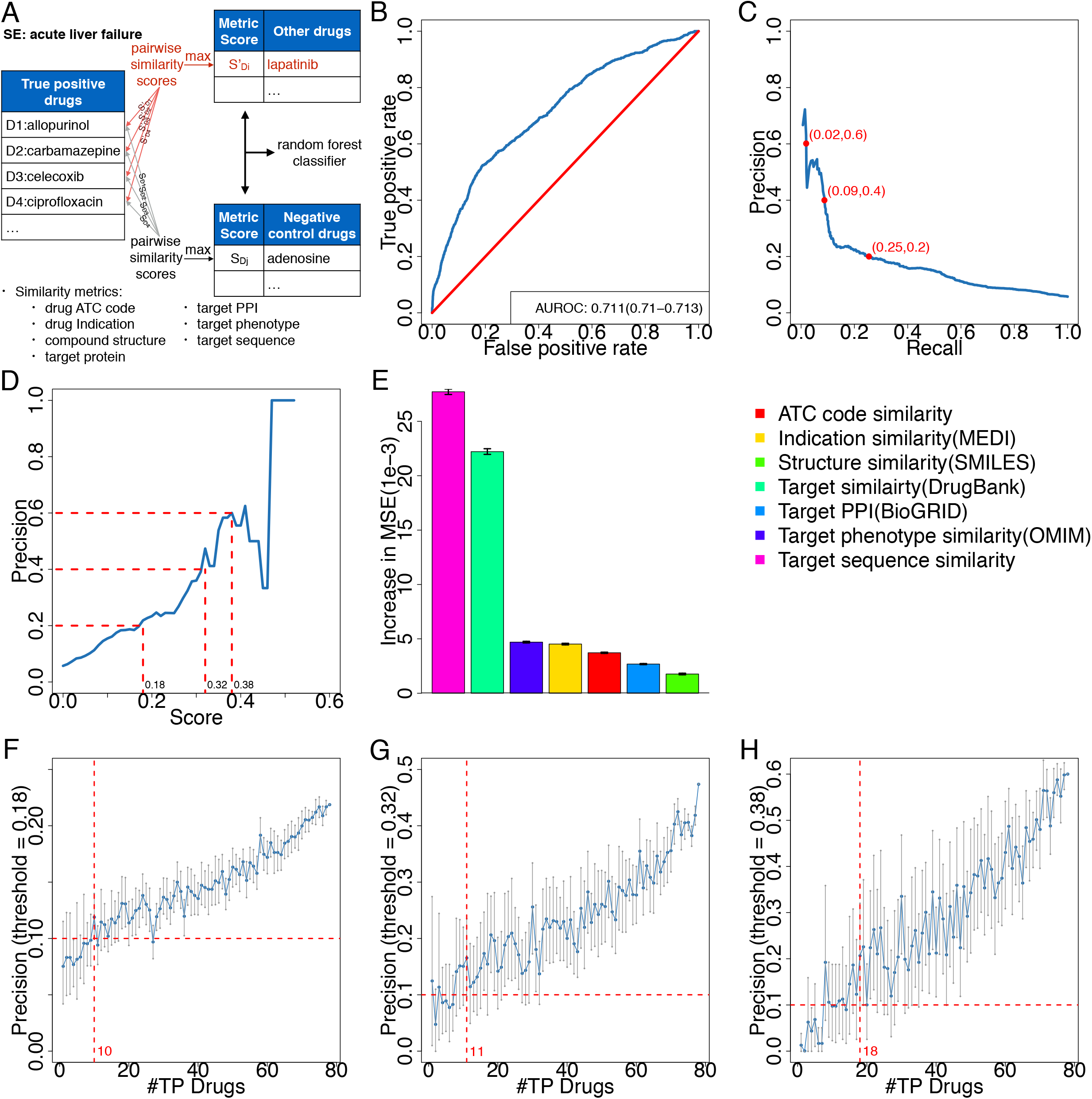
Performance of SIDERctrl model. **(A)**: A brief workflow of SIDERctrl. Each unreported drug (drugs that are not annotated with the side effect) was scored by the max similarity to all true positive drugs in seven features. Then a random forest classifier was built to differentiate negative control drugs from other unreported drugs. **(B)**: Receiver operating characteristic (ROC) curve (blue) of SIDERctrl. Average performance of 100 run was plotted to account for the stochastic nature of random forest. The area under ROC curve (AUROC) value and 95% confidence interval were annotated at the bottom right of the plot. A random classifier will have ROC curve as diagonal (red) and AUROC value of 0.5. **(C)**: Precision-recall curve (blue) of SIDERctrl. Precision = TP/(TP+FP), Recall = TP/(TP+FN). Average performance of 100 run was plotted. Precision decreases from ~0.7 to ~0.2 when recall increases from 0 to 0.1, then decreases in a lower rate to 0.05 as recall increases to 1. Three thresholds of precision (0.2, 0.4, and 0.6) and the corresponding recall value were annotated in red on the curve. **(D)**: Precision-predicted score curve (blue) of SIDERctrl. Dashed lines (red) show the three thresholds of score (0.18, 0.32, and 0.38) we defined, in order to reach minimum precision of 0.2, 0.4, and 0.6 on the training set, respectively. **(E)**: Barplot showing the predictive power of seven features in SIDERctrl, which was measured by the increase in mean squared error (MSE) when the feature was removed from the model. The normalized value was shown in the magnitude of 1.0E-3. Error bar shows 95% confidence interval of the average. All seven features were ranked from highest to lowest. Description of each feature was shown in the figure legend on the right. **(F-H)**: Line charts (blue) showing the relationship between number of true positive (TP) drugs and the resulting precision of SIDERctrl. Each point shows the average precision (y-axis) of models built by a particular number of TP drugs (x-axis). Error bars (grey) show the 95% confidence interval of the average. Precision was calculated under three thresholds: 0.18 **(F)**, 0.32 **(G)**, and 0.38 **(H)**. Dashed lines (red) show the minimum number of TP drugs required (10,11, and 18) to reach a minimum precision of 0.1.

### 2.4 Estimate the performance of new predictions by sampling subsets from the training set

We investigated the association between the number of true positive drugs and model performance by sampling subsets of true positive drugs from the training set to recalculate the similarity scores of unreported drugs. A new SIDERctrl model was built based on the scores, and evaluated by the same method described above. To account for the stochastic nature of randomization, the analysis was repeated 20 times for every possible size of the subset. The average result of each size was used to estimate the performance of new predictions made for SEs with the same number of true positive drugs. Using this method, we were able to make estimations for SEs annotated with fewer true positive drugs than the training set. For SEs with more true positive drugs, we gave a minimum estimation as the performance derived from the whole training set.

### 2.5 Validate the predicted negative controls using data from AEOLUS

AEOLUS dataset was downloaded from Dryad [5], and detailed in Supplementary Methods. Drugs and SEs were matched between SIDER and AEOLUS using the concept name from the “concept” table. Relationship between drugs and SEs were extracted using the proportion reporting ratio (PRR) statistics from the “standard_drug_outcome_statistics” table. A drug was defined as true positive of a SE if the lower confidence interval of PRR score is greater than 1.

## 3 RESULTS

### 3.1 SIDERctrl achieved a 42% improvement over random classifier

As described above, SIDERctrl was trained on a dataset that contains 1,824 relationships between four SEs and 456 drugs, including 78 positive, 100 negative, and 1646 other relationships. Overall, SIDERctrl achieved an AUROC of 0.711 (95% CI: 0.710-0.713; Fig. 1B), increasing the performance of random classifier by 42%. The precision of SIDERctrl reached 0.2, 0.4, 0.6 when unreported drugs with predicted score greater than 0.18, 0.32, 0.38 were defined as negative controls, respectively (Fig. 1C,D). SIDERctrl incorporated seven pharmacological features in the random forest model, where each unreported drug was scored by the maximum feature similarity to true positive drugs. We compared the predictive power of distinct features. Target sequence similarity and target similarity were found to have the highest power among seven features (Fig. 1E).

### 3.2 The performance of SIDERctrl is positively correlated with the number of true positive drugs

We found that the average precision of SIDERctrl increases linearly while the variation decreases as the number of true positive drugs used to generate the feature scores of unreported drugs increases from 1 to 78 (Fig. 1F-H). Similar results were observed in the other measurements of performance such as AUROC (Supplementary Fig. 1A) and recall (Supplementary Fig. 1B-D). The precision of SIDERctrl remains consistently greater than 0.1 (*P* < 0.05) when more than 10, 11, 18 true positive drugs were used to generate the similarity scores of unreported drugs under the threshold of 0.2, 0.4, 0.6, respectively. Therefore, we set the minimum number of true positive drugs required for the SIDERctrl model as 10, 11, and 18 under three thresholds, in order to guarantee a minimum precision of 0.1 in the predicted negative controls.

### 3.3 Predicted negative controls exhibit lower false negative rate compared to other unreported drugs

We then applied SIDERctrl to predict negative controls for 890 SEs in SIDER annotated with more than 10 true positive drugs (Supplementary Methods). Using the thresholds defined above, we obtained three sets of predicted negative controls (Table 1; Supplementary Table 1). The precision of our predictions was estimated to be at least 0.15, 0.26, and 0.37 in three sets. The number of predicted drugs dropped significantly as higher thresholds were applied. On average, 29.4±0.8, 4.2±0.2, 2.0±0.1 drugs were predicted per SE under three thresholds. No SE was predicted with more than 10 drugs under the highest threshold of 0.38.

**Table 1:** summary of our predictions for 890 SEs in SIDER. SEs: side effects; TP: true positive drugs; PN: predicted negative controls.

| Threshold of score | 0.18 | 0.32 | 0.38 |
| --- | --- | --- | --- |
| Minimum #TP required | 10 | 11 | 18 |
| #Predicted SEs | 890 | 772 | 458 |
| #SEs with $\geq 10$ PN | 845 | 36 | 0 |
| #True positive drugs | 62.7(57.8-67.8) | 51.8(48-55.3) | 62.6(57.9-67.1) |
| #Predicted negative controls | 29.4(28.6-30.2) | 4.2(4.0-4.4) | 2.0(1.9-2.1) |
| Estimated precision (%) | $\geq 15$ | $\geq 26$ | $\geq 37$ |

We validated our predictions by comparing the false negative rate between predicted negative controls and other unreported drugs, which was defined as the proportion of true positives reported in an independent resource. More than 97% of the 890 SEs can be validated using this approach. The false negative rates of predicted negative controls are 6.1±0.4%, 5.6±1.0%, and 7.2±2.1% under three thresholds (Fig. 2A), significantly lower than those of other unreported drugs: 9.0±0.3% (*P* = 2.3e-33), 8.9±0.3% (*P* = 4.9e-10), and 9.3±0.4% (*P* = 2.7e-2). We also grouped the 890 SEs into 19 categories based on human body system (Supplementary Methods), and observed similar results across distinct systems (Fig. 2B-F & Supplementary Fig. 2A-N). In 12 of the 19 systems, at least one of three predicted sets exhibits lower false negative rate than other unreported drugs.

**Fig. 2:**
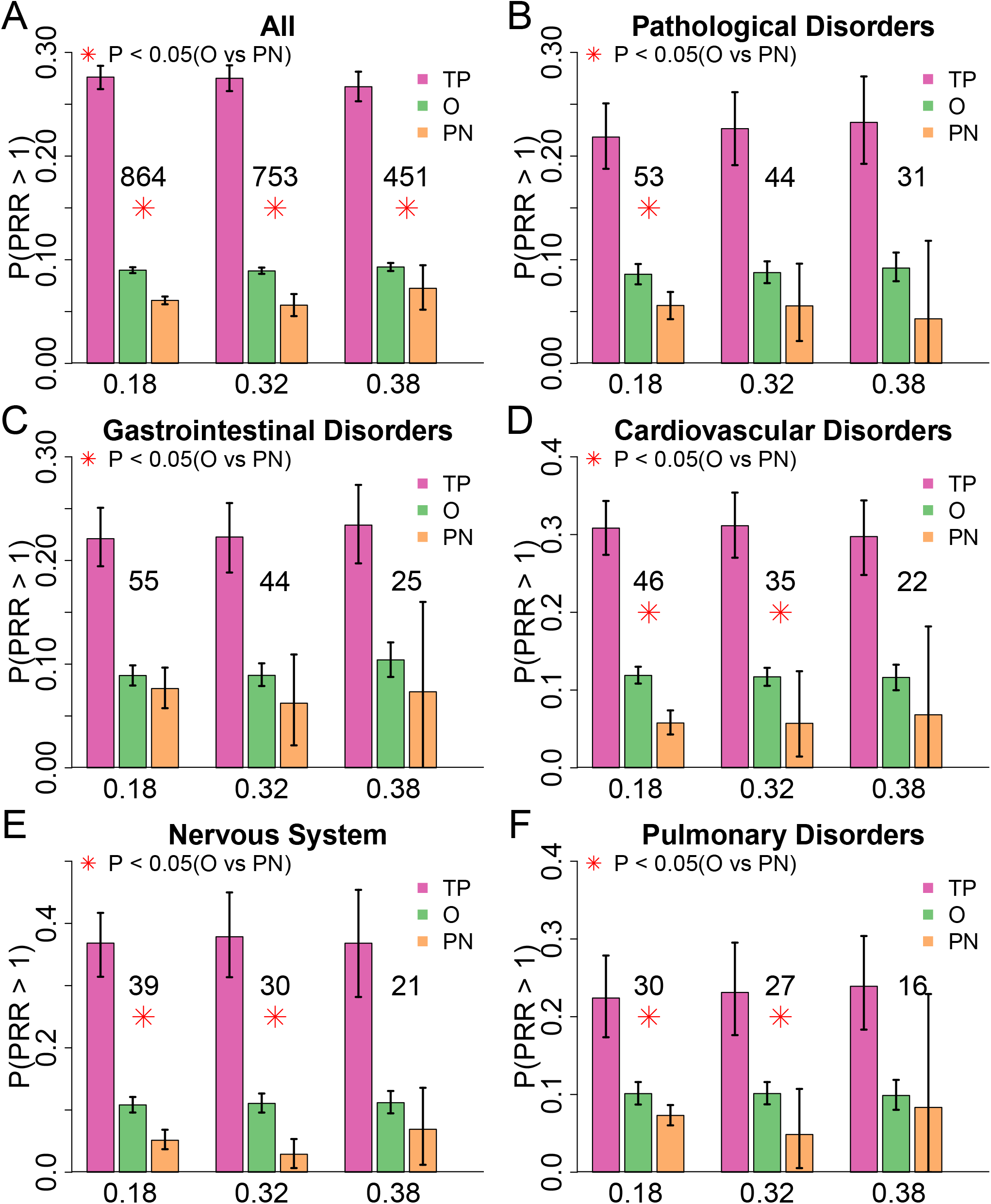
Validation of SIDERctrl predictions using AEOLUS data. TP: true positive drugs; 0: other drugs; PN: predicted negative controls. **(A)**: Barplot showing the false negative rate (y-axis) in TP (pink), 0 (green), and PN (orange). The false negative rate was defined as the proportion of drugs with proportional reporting ratio (PRR) score greater than 1 from AEOLUS. Results under three thresholds (0.18, 0.32, and 0.38) were shown in the plot. Each bar shows the average value of all validated side effects (SEs) while the error bar shows 95% confidence interval of the average. The number of validated SEs was shown in the plot. A red star (*) was shown under the number when the false negative rate of PN is significantly lower than that of 0 (*P* < 0.05). **(B-F)**: All the validated SEs shown in (A) were grouped into **19** categories based on human body systems. **(B-F)** show the five systems with highest number of SEs. The name of each system was shown as the title. Results of the rest 14 systems can be found in Supplementary Fig. 2.

### 3.4 Predicted negative controls of similar SEs show higher variability than true positive drugs

We investigated whether similar SEs were predicted with similar negative controls by comparing the Jaccard similarity of predictions within the same system of SEs and across different systems. We only found a significant difference between the two groups when using 0.18 as the threshold to define negative controls (P = 0.045; Fig. 3A), while no significant different was detected between the two groups under higher thresholds (Fig. 3B,C). As comparison, the same analysis was performed on the true positive drugs that were used to generate the feature scores. We found that the within-system similarity is consistently higher than the across-system similarity under all three thresholds (Fig. 3A-C), proving that similar SEs were annotated with similar sets of true positive drugs in SIDER. Altogether, the results show that similar sets of true positive drugs may result in substantial difference among the predicted negative controls.

**Fig. 3:**
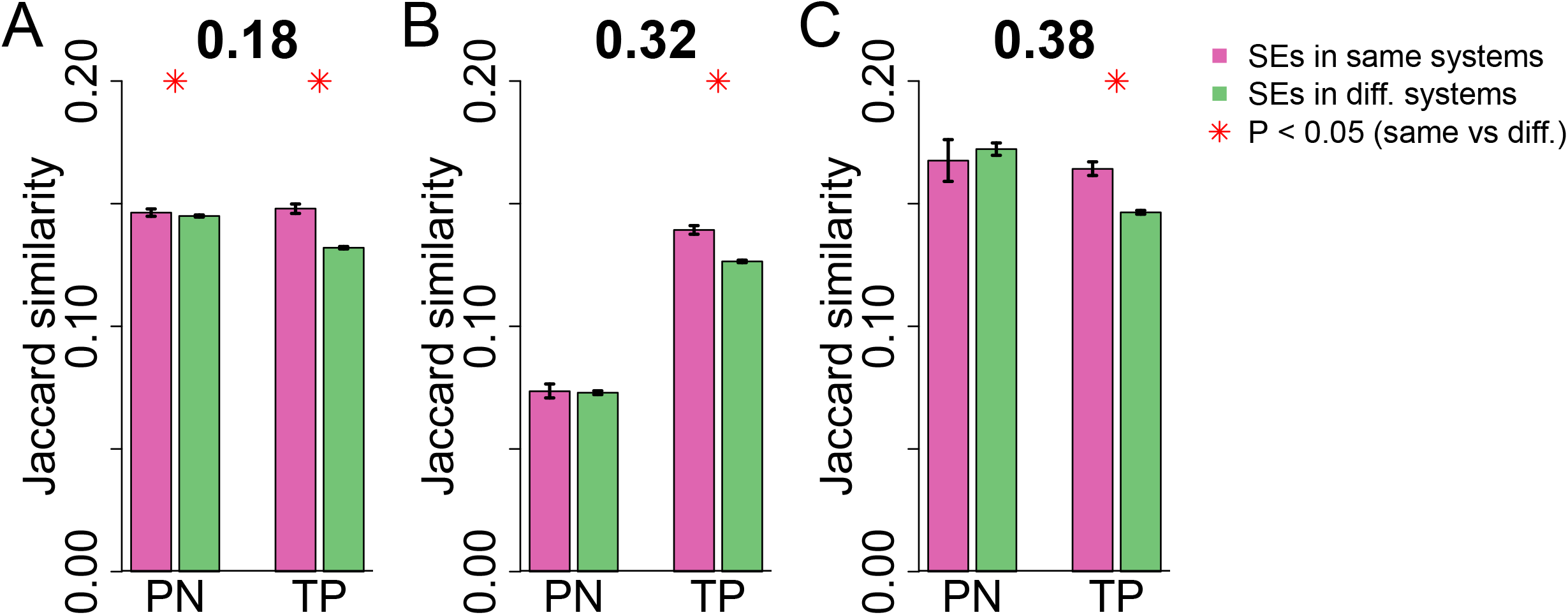
Comparison of similarity in SIDERctrl predictions among distinct side effects. PN: predicted negative controls; TP: true positive drugs. **(A-C):** Barplot showing the similarity of TP and PN among SEs within the same categorical system (pink) and across different systems (green). Similarity was measured by the Jaccard similarity (#intersect/#union) between two sets of drugs (y-axis). Each bar shows the average pairwise similarity of all possible pairs while the error bar shows 95% confidence interval of the average. A red star (*) was shown above the bars when the within-system similarity is significantly higher than the across-system similarity (*P* < 0.05). Results under three thresholds were shown: 0.18 **(A)**, 0.32 **(B)**, and 0.38 **(C)**. Results by each categorical system can be found in Supplementary Fig. 3.

## 4. DISCUSSION

One important issue in drug safety research is the lack of high-qualify reference set. Despite the wide use of reporting systems such as FAERS, many SEs of drugs remain unreported. Such incompleteness causes a high false negative rate among all unreported drugs. In this paper, we introduced a supervised learning model that addressed this issue by ranking all unreported drugs based on their similarity to the reported positive drugs. Drugs with lower similarity were predicted as negative controls. We set three different thresholds of predicted score to define negative controls, in order to provide users with more choices. Higher thresholds result in fewer, but more accurate predictions. We also found that when more true positive drugs were reported and used to generate the similarity scores of unreported drugs, better performance of SIDERctrl was obtained. This is because more information can be learned by our model regarding the chemical and biological properties of drugs associated with the SE.

The highlight of this study is that we provided three sets of predicted negative controls for all 890 SEs in SIDER that were annotated with more than 10 true positive drugs. We validated our predictions using data from an independent resource AEOLUS, where we showed that the false negative rate of predicted negative controls is about two-thirds of other unreported drugs. In addition, we found that while similar SEs may share similar true positive drugs, they are less likely to share predicted negative controls. The high variability in the predicted negative controls matches a finding described earlier in the paper that minor change in the true positives of training set leads to variation in performance of SIDERctrl (Fig. 1F-H & Supplementary Fig. 1 A-D). Among the three sets, the ones generated by thresholds of 0.32 and 0.38 show higher precision, but contain very few predictions for most SEs. In those cases, the few predicted drugs are not enough for research purposes such as building supervised learning models to predict the occurrence of SEs. However, we think the high-quality sets may be useful in the study of experimental toxicology. In such scenario, SIDERctrl serves as a preliminary screen of negative controls to be used in the experiment. Researchers can apply the predicted drugs to cell line assays or organ-on-a-chip systems, and compare the results with those derived from true positive drugs to study the cellular mechanism of side effects and identify related off-target proteins. The set generated by threshold of 0.18, on the other hand, contains much more predictions per SE, thus can be used as reference set for predicting the occurrence of SEs or drug-induced toxicity in human body systems and tissues. We expect the new resource to improve the data quality in drug safety research and facilitate the generation of new hypotheses studying the mechanism of SEs.

## AUTHORS CONTRIBUTIONS

Yun Hao and Nicholas P. Tatonetti designed the research. Yun Hao performed the research and analyzed the data. Yun Hao and Nicholas P. Tatonetti wrote the manuscript.

